# Structural Mechanism of Electron Shuttling in Inducible Nitric Oxide Synthase

**DOI:** 10.64898/2026.04.17.719136

**Authors:** Yiting Shi, Xiaoyu Liu, Lei Chen

## Abstract

Nitric oxide (NO) signaling is pivotal in numerous physiological processes and is implicated in a spectrum of human diseases. Nitric oxide synthases (NOS) initiate NO signaling and govern its magnitude and duration, making them key drug targets. Despite decades of investigation, the structural mechanism by which NOS enzymes transfer electrons from NADPH to haem remains incompletely understood. Here, we report cryo-electron microscopy (cryo-EM) studies of the inducible NOS (iNOS) homodimer in complex with calmodulin (CaM) captured under catalytic turnover conditions, resolving two important functional states: the electron input state and output state. In the input state, the FMN-binding subdomain (FMND) docks onto the FAD/NADPH-binding subdomain (FNR), positioning the FMN cofactor to accept electrons from FAD. The FMND then undergoes a large rotational movement to engage the oxygenase domain (OXY) of the other protomer, adopting the output state, which enables electron transfer from FMN to the haem center via W366. This dynamic movement of the FMND shuttles electrons from the reductase domain (RED) to the OXY active site in iNOS. A point mutation (S594E) that disrupts the FMND–OXY interface markedly reduces the catalytic activity of iNOS and traps a fraction of enzymes in a non-productive intermediate conformation. Together, these findings elucidate the structural mechanism of FMND-mediated electron transfer in the iNOS catalytic cycle.

## Introduction

Nitric oxide (NO) is a unique gaseous signaling molecule that plays an essential role in cardiovascular homeostasis, cellular metabolism, neurotransmission, and immune defense ^1^. NO signaling is initiated by NO production via nitric oxide synthase (NOS) (Fig. 1a); NO then freely diffuses to target cells, where it either covalently modifies macromolecules or binds to its primary receptor, soluble guanylate cyclase (sGC), to generate the second messenger cGMP, which activates downstream signaling cascades ^1,2^. As such, NOS activity is pivotal to several key physiological processes, including vasodilation and blood pressure regulation, synaptic plasticity and neuronal communication, and pathogen clearance by immune cells ^1,3^. Overproduction of NO correlates with the hypotension and tissue damage observed in septic shock ^4^, as well as pancreatic β-cell malfunction in diabetes mellitus ^5^. Underproduction of NO is implicated in endothelial dysfunction, contributing to hypertension and atherosclerosis ^3^. Therefore, NOS are important drug targets ^6,7^.

**Figure 1.**
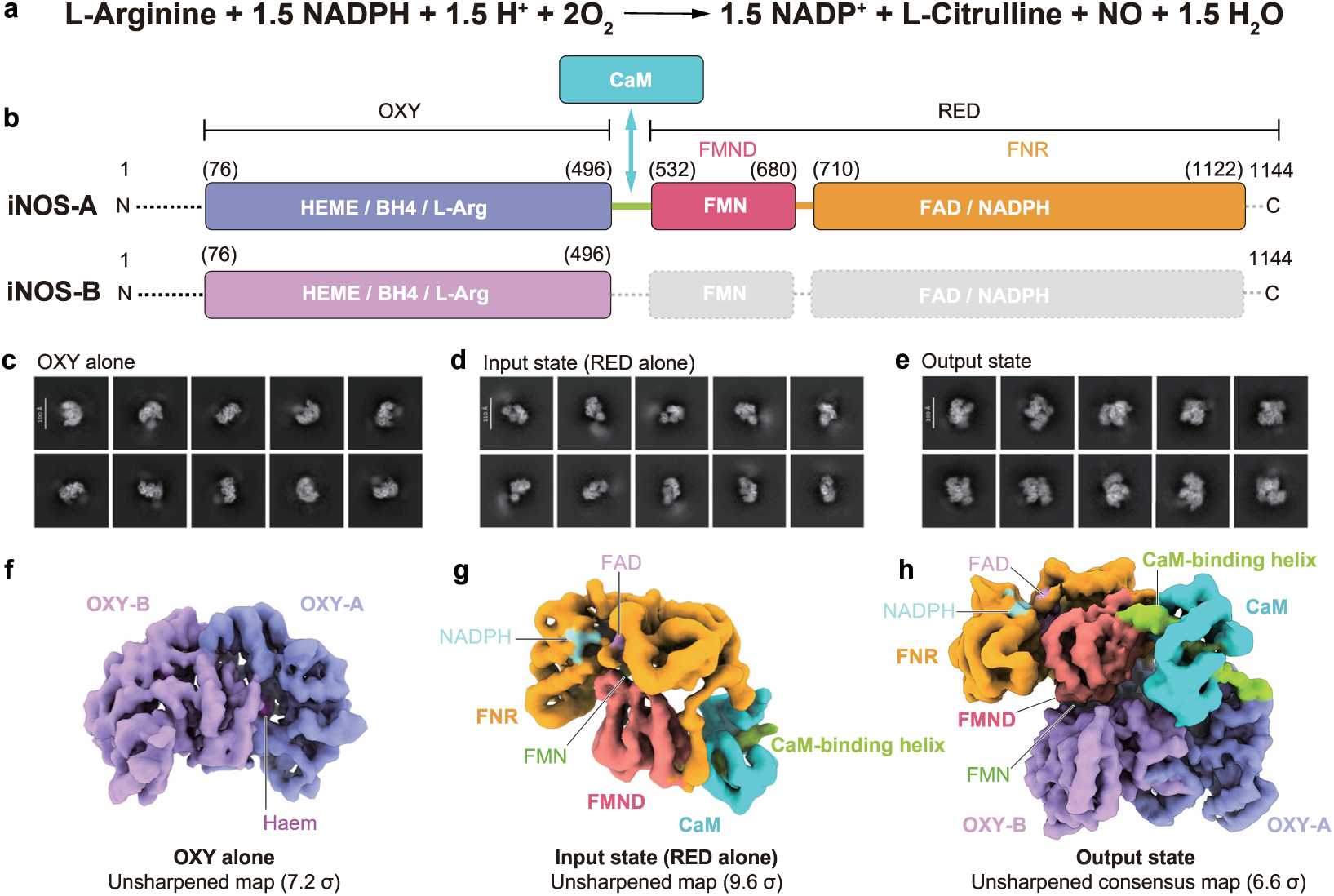
Structure of iNOS under the turnover condition. a, The chemical reaction catalyzed by iNOS. b, The domain composition of the mouse iNOS holoenzyme. Structured domains are shown as boxes, while flexible regions invisible in cryo-EM structures are indicated by dashed lines. The iNOS protomer with well-resolved density of the RED and bound CaM is designated as iNOS-A, while the other iNOS protomer with only recognizable density of the OXY is referred to as iNOS-B. The OXY, CaM-binding helix, FMND, and FNR of iNOS-A are colored purple, light green, salmon, and orange, respectively. The OXY of iNOS-B is colored light purple. Regions not included in the final model are shown as dashed lines or boxes. Brackets above indicate domain boundaries (middle) and designate visible regions (at each terminus). CaM is colored cyan, and the arrow pointing toward the CaM-binding helix indicates their interaction. c, 2D classes representing the OXY dimer alone. d, 2D classes representing the putative input state (RED alone), with clear features of both the FNR and FMND. e, 2D classes representing the putative output state, with both the RED and OXY dimer. f, Unsharpened map of isolated iNOS OXY dimer colored as in b. g, Unsharpened map of iNOS RED in the input state colored as in b. h, Unsharpened consensus map of iNOS RED in the output state colored as in b.

Three isoforms of NOS exist in humans: inducible NOS (iNOS), neuronal NOS (nNOS), and endothelial NOS (eNOS) ^8^, and their activities are highly regulated. iNOS is expressed in macrophages and various other cell types, with its transcription initiated only upon immune stimulation ^9^. iNOS binds calmodulin (CaM) tightly ^10^ and is constitutively active independent of calcium signaling ^11,12^, enabling sustained NO production. In contrast, nNOS and eNOS are activated upon CaM binding only in the presence of elevated intracellular calcium ^13^, and their activities are thus tightly regulated by calcium signaling. Genetic variations in iNOS are associated with asthma ^14^; mutations in the gene encoding nNOS lead to hypogonadotropic hypogonadism ^15^; and polymorphisms in the eNOS gene are associated with cardiovascular diseases ^16^, highlighting the physiological importance of NOS enzymes.

NOS are NADPH-dependent cytochrome P450 oxidoreductase-like enzymes ^17^ that encompass an N-terminal haem-containing oxygenase domain (OXY) and a C-terminal reductase domain (RED), with a CaM-binding domain in the middle ^13^. The RED can be further divided into an FMN-binding domain (FMND) and an FNR-like FAD/NADPH-binding domain (FNR) ^17^ (Fig. 1b). Previous biochemical and biophysical studies have established the electron transfer pathway in NOS ^17^. Upon NOS activation, NADPH transfers a hydride to FAD within the FNR; electrons are then transferred from FAD to the FMN bound in the FMND (electron input to FMND) and subsequently to the haem in the OXY (electron output from FMND) ^17^. The OXY utilizes electrons to activate oxygen for arginine oxidation and NO production in a H4B-dependent manner ^18^. Structural studies on individual domains of NOS enzymes have uncovered the zinc-mediated dimeric architecture of the OXY ^19,20^, the mechanism of CaM binding ^21,22^, and the configuration of the RED in the input state of nNOS ^23–25^. However, in the input state structure, the FMN cofactor bound in the FMND is shielded by the FNR and is not accessible to the OXY ^23^. Therefore, a major conformational change of the RED is required for electron output from FMN. Moreover, the FMND must dock onto the OXY in a specific manner to allow efficient electron transfer to haem.

Studies using hydrogen-deuterium exchange ^26^, negative stain electron microscopy ^27,28^, and crosslinking mass spectrometry ^29,30^ have demonstrated that the FMND can transiently dock onto the OXY to adopt an output state that is competent for efficient electron transfer to the haem cofactor. More recently, an output state-like conformation has been visualized at low resolution in the cyanobacterial syNOS protein via cryo-EM ^31^. However, the detailed molecular mechanism by which the FMND reorients and docks onto the OXY domain—particularly at high resolution and within the context of the full-length mammalian holoenzyme—has remained elusive. Here, we report cryo-EM structures of wild-type mouse iNOS captured under turnover conditions, which reveal distinct conformations of the enzyme corresponding to both the input state (FMND docked onto FNR) and the output state (FMND docked onto OXY). We also present the structure of a loss-of-function mutant, S594E, some particles of which are trapped in a non-productive intermediate state. Together, these structures provide a comprehensive view of the structural mechanism governing FMND-mediated electron shuttling in iNOS.

## Results

### Structure determination of iNOS under turnover conditions

To better mimic the cytosolic environment of eukaryotic cells, we co-expressed mouse iNOS and CaM in Expi293F mammalian cells using the BacMam system ^32,33^. The iNOS holoenzyme expressed in this manner exhibited robust NO-producing activity (Extended Data Fig. 1a). Using purified iNOS protein (Extended Data Fig. 1b-c), we prepared cryo-EM samples under aerobic conditions in the presence of all cofactors and substrates, including NADPH, FAD, FMN, BH4, and arginine, thereby mimicking the catalytic conditions of iNOS (Fig. 1 and Extended Data Fig. 2). During the extensive 2D classification process, we observed class averages corresponding to the isolated OXY dimer of iNOS (Fig. 1c) and the isolated RED (Fig. 1d). We also identified several 2D classes in which prominent additional densities appeared beside the OXY dimer (Fig. 1e), suggesting the binding of the RED to the OXY dimer.

Subsequent 3D reconstruction and classification resolved three major classes (Fig. 1f-h and Extended Data Fig. 2). In the first 3D class, we observed an isolated OXY dimer (Fig. 1f), with a resolution of 3.5 Å (Extended Data Fig. 2 and Supplementary Table 1). In the second 3D class, we observed an isolated RED in complex with CaM, detached from the OXY dimer (Fig. 1g, Extended Data Fig. 3 and Supplementary Table 1). Moreover, the RED adopts a conformation in which NADPH is ready to transfer a hydride to FMN (Fig. 2). We refer to this structure as the input state, which was refined to a resolution of 3.8 Å (Extended Data Fig. 3 and Supplementary Table 1). In the third 3D class (Fig. 1h), the RED and its associated CaM from one subunit bind to the OXY of the other subunit, adopting a conformation poised for electron transfer from FMN to haem (Fig. 3). We refer to this structure as the output state, which was refined to a resolution of 3.4 Å (Extended Data Fig. 4-8 and Supplementary Table 1). The observation of both the input and output states in the same sample is consistent with a model in which NOS interconverts between these two states to transfer electrons from NADPH bound in the FNR to haem bound in the OXY under turnover conditions.

**Figure 2.**
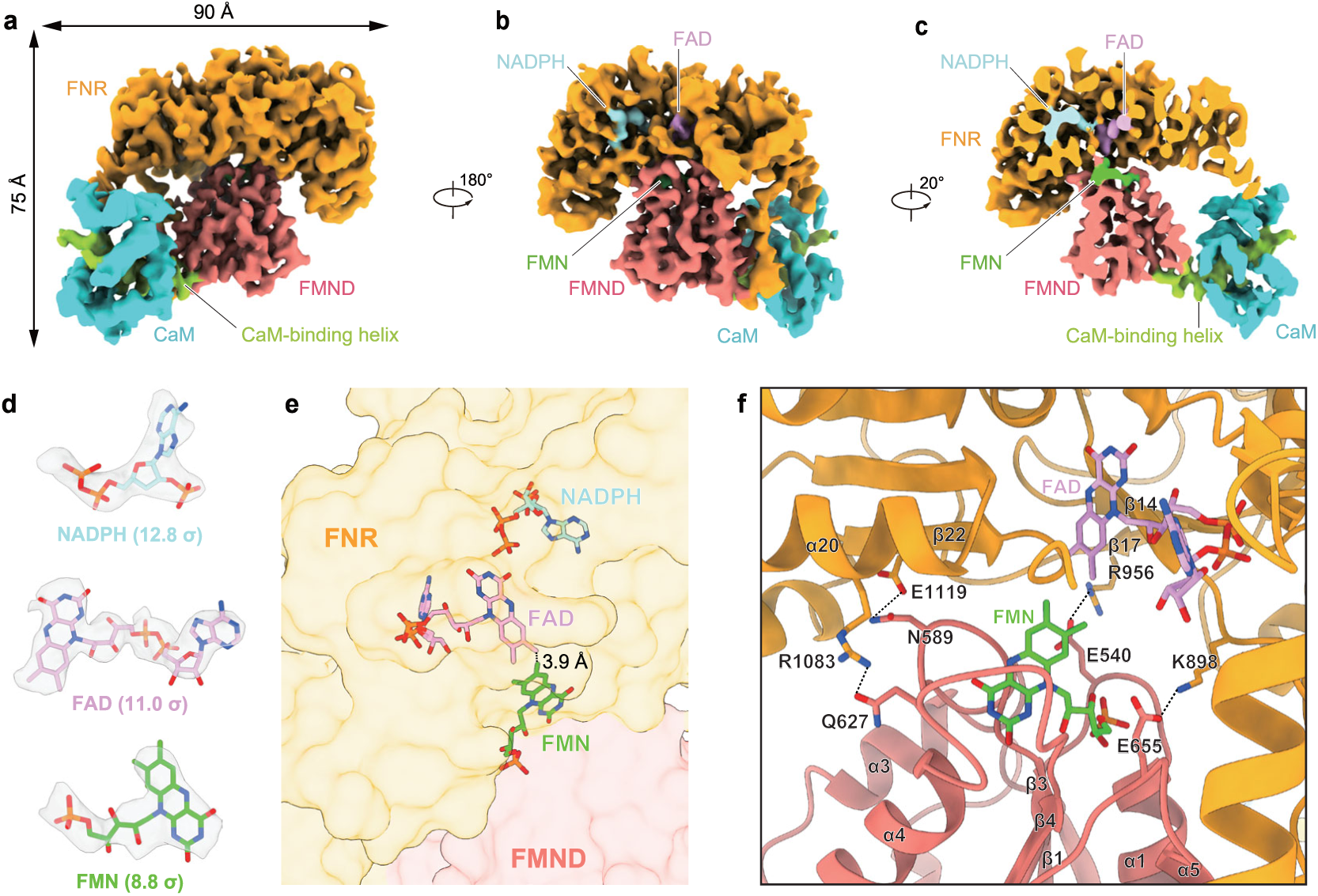
Structure of iNOS RED-CaM complex in the input state. a, Cryo-EM map of iNOS in the input state. Each subunit is colored as in Fig. 1b. b, A 180°-rotated view of a. c, Cut-open view of the cryo-EM map of iNOS RED-CaM complex in the input state. FMN, FAD and NADPH are colored green, hot pink and light blue, respectively. d, Densities of substrates and cofactors found in the iNOS RED-CaM complex in the input state. Densities corresponding to NADPH, FAD, and FMN were contoured at 12.8 σ, 11.0 σ, and 8.8 σ, respectively. The nicotinamide group of NADPH is not modeled due to its poor density. e, Edge-to-edge distance between FAD and FMN in the input state is shown beside the dashed line. Ligands are shown as sticks, and iNOS is shown as a semi-transparent surface. f, The interactions between FMND and FNR of iNOS RED in the input state.

**Figure 3.**
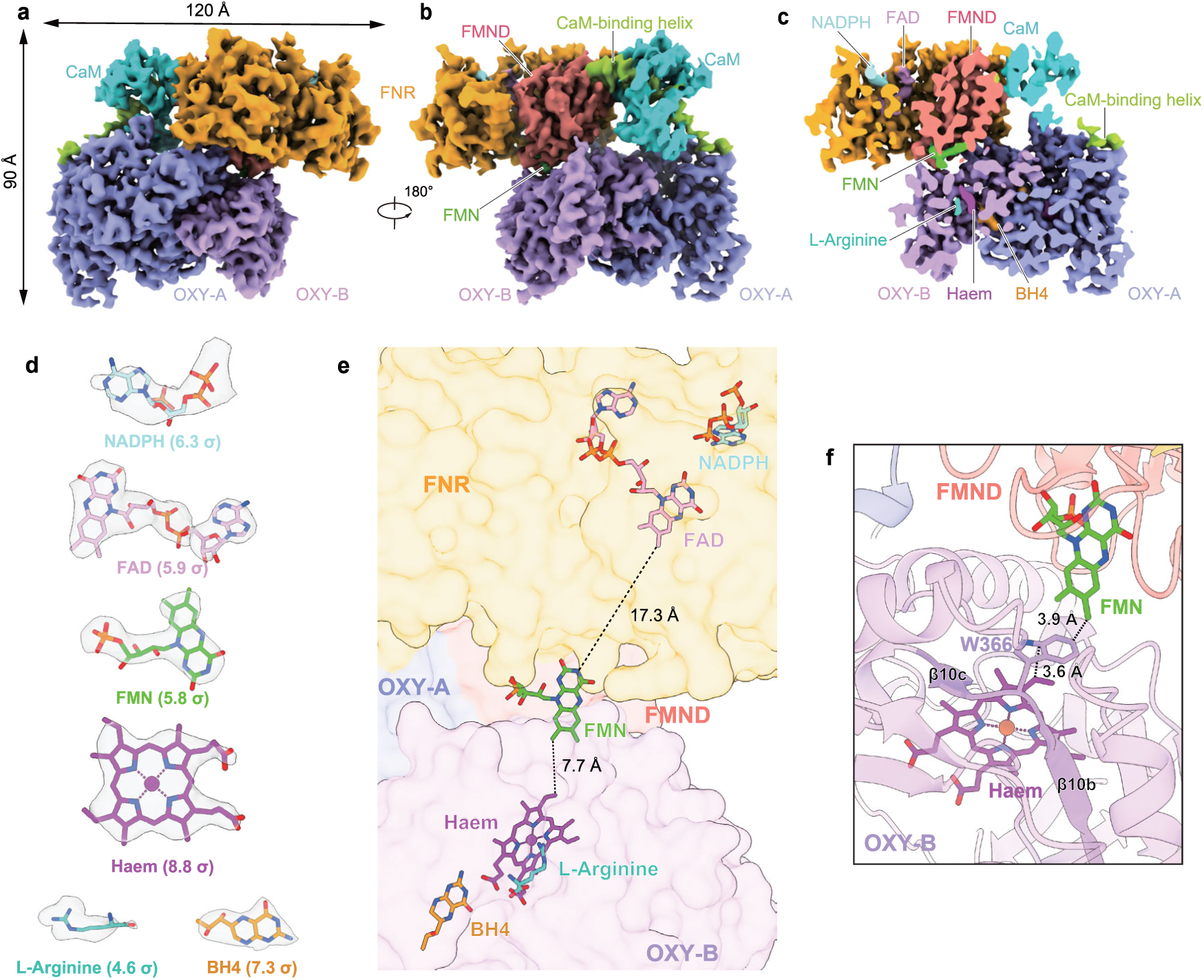
Structure of iNOS in the output state. a, Side view of the cryo-EM map of the iNOS-CaM complex in the output state. Each subunit is colored as in Fig. 1b. b, A 180°-rotated view of a. FMN is colored green. c, Cut-open view of b. Haem, BH4, L-Arginine, FAD, and NADPH are colored magenta, bright orange, sea blue, hot pink, and light blue, respectively. d, Densities of substrates and cofactors found in the iNOS-CaM complex in the output state. Densities corresponding to NADPH, FAD, FMN, Haem, L-Arginine, and BH4 were contoured at 6.3 σ, 5.9 σ, 5.8 σ, 8.8 σ, 4.6 σ, and 7.3 σ, respectively. The nicotinamide group of NADPH is not modeled due to its poor density. e, Relative position of cofactors and substrates in the output state structure. Edge-to-edge distance between the isoalloxazine groups of FMN and FAD, as well as that between FMN and haem, are denoted beside the dashed lines. f, Zoom-in view of the configuration between FMN, W366, and haem in the output state. Edge-to-edge distances between FMN and W366 and between W366 and haem are denoted beside the dashed lines.

### Structure of the RED-CaM complex in the input state

In the electron density map of the input state (Fig. 2a-c), we observed clear densities representing FAD, FMN, as well as the adenine–ribose–phosphate portion of NADPH (Fig. 2d). Notably, the density for the nicotinamide group of NADPH is poor (Fig. 2d), suggesting its high flexibility during catalysis; therefore, this group was not modeled. In this structure, the isoalloxazine group of FMN is in close proximity to that of FAD, with an edge-to-edge distance of 3.9 Å (Fig. 2e)—well within the range for optimal electron transfer. However, because the FMND is detached from the OXY dimer in the input state, direct electron transfer between FMN and haem is not favorable.

We observed multiple polar interactions between the FMND and FNR (Fig. 2f). E540 on the β1-α1 loop of the FMND forms an electrostatic interaction with R956 on β17 of the FNR (Fig. 2f). Previous studies have shown that mutations of either of these two residues in nNOS (E762 and R1229) decrease NO production ^34^. E655 on the β5-α5 loop of the FMND interacts with K898 on the α13-β14 loop of the FNR (Fig. 2f). N589 on the β3-α3 loop of the FMND forms a hydrogen bond with E1119 on β22 of the FNR (Fig. 2f). Q627 on the β4-α4 loop of the FMND forms a hydrogen bond with R1083 on α20 of the FNR (Fig. 2f). These interactions hold the FMND in the FNR at an optimal pose for electron input. Notably, when comparing this structure with that of the nNOS RED (PDB ID: 1TLL) ^23^, we observed a 25°rotation of the FMND calculated using Dyndom 3D ^35^ (Extended Data Fig. 5a). This difference may be attributed to intrinsic structural differences between nNOS and iNOS or the binding of CaM. Moreover, some interaction pairs in the input state are conserved between nNOS and iNOS, including E540-R956 and N589-E1119 (Extended Data Fig. 7-8).

### Redox center configuration of iNOS in the output state

In the electron density map of the output state (Fig. 3), we observed clear densities representing FAD, FMN, haem, BH₄, arginine, and the adenine–ribose–phosphate portion of NADPH (Fig. 3d). The positions of haem, BH4, and arginine are consistent with those observed previously in the isolated OXY dimer (PDB ID: 1NOD) ^36^ (Extended Data Fig. 4i-k). Moreover, the isoalloxazine group of FMN is far from that of FAD (Fig. 3e), with an edge-to-edge distance of 17.3 Å (Fig. 3e), suggesting that direct electron transfer from FAD to FMN is unfavorable. In contrast, FMN is closer to haem bound in the OXY of the other subunit, with an edge-to-edge distance of 7.7 Å (Fig. 3e). More importantly, FMN is near W366 on the β10b-β10c loop in the OXY (Fig. 3f), with an edge-to-edge distance of 3.9 Å (Fig. 3f). W366 is close to the haem, with an edge-to-edge distance of 3.6 Å (Fig. 3f). This configuration of redox centers enables efficient electron transfer from FMN to haem via W366 in the output state and is consistent with previous studies suggesting that W366 participates in the electron transfer process from FMN to haem ^26^.

### Interdomain interactions of iNOS in the output state

In the output state structure, only one RED is docked onto the OXY dimer; the other RED is not visible, likely due to its high flexibility (Fig. 3a-b and Fig. 4a). Calcium-bound CaM wraps around the CaM-binding helix of iNOS (Fig. 4b). The electron density of CaM is blurry and poorer than that of the rest of the complex, suggesting greater mobility of CaM (Fig. 3b). This observation is consistent with the lack of a stable interface between CaM and the OXY dimer in the output structure. We also did not observe a stable interface between the FMND and the FNR in this structure. Instead, we identified an interface between the FNR and OXY and an interface between the FMND and OXY (Fig. 4c-e).

**Figure 4.**
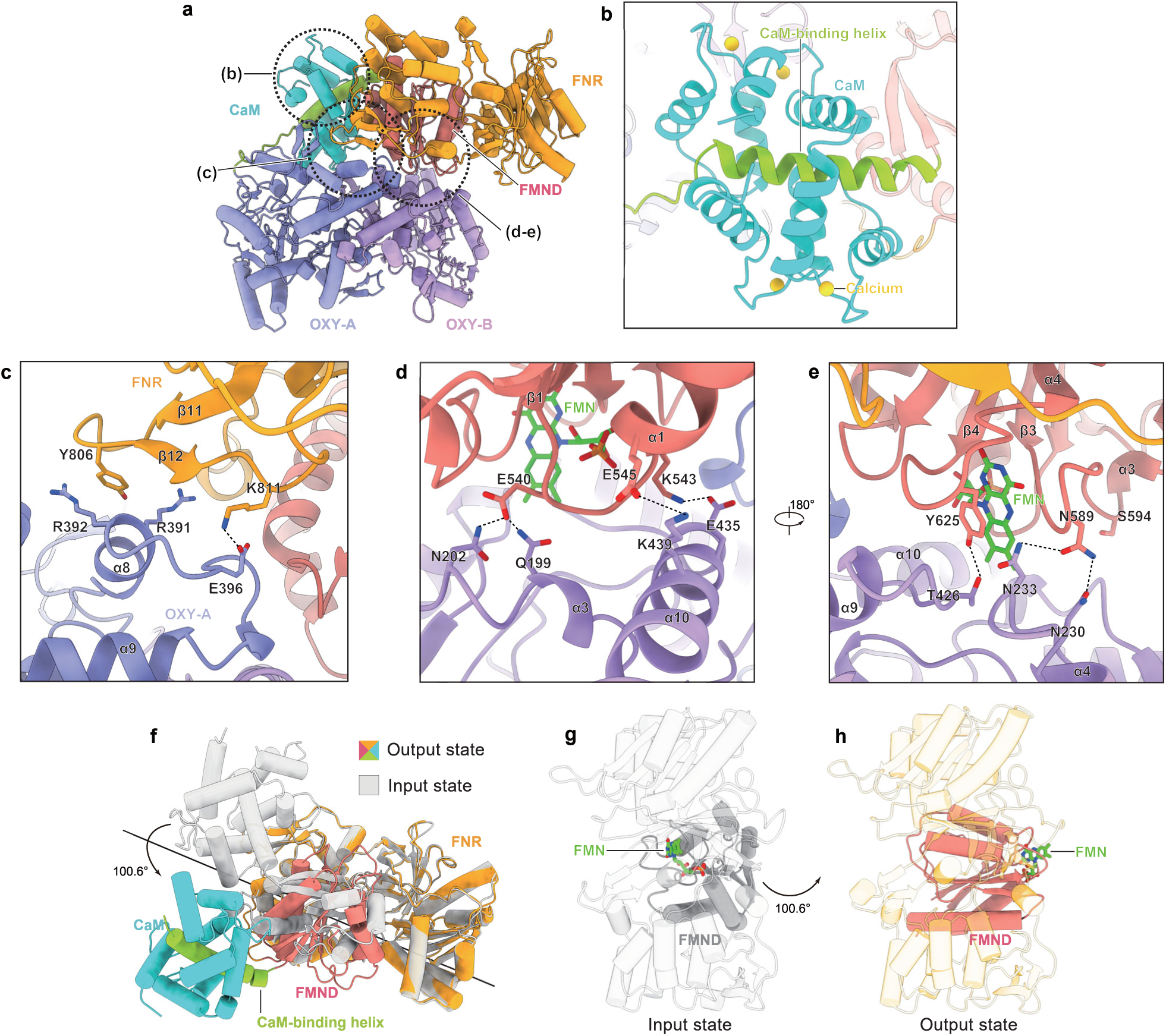
Interdomain interactions of iNOS in the output state. a, Side view of the iNOS structure in the output state, with helices shown as cylinders. b, Binding between calcium-bound CaM and the CaM-binding helix of iNOS (circled in a). c, Interface between iNOS-A FNR and OXY-A. The putative salt bridge is shown as dashes. d, Interface between iNOS-A FMND and OXY-B. Putative salt bridges and hydrogen bonds are shown as dashes. The position of S594 on the FMND is also indicated. e, A 180°rotated-view of d. f, Structural comparison between iNOS in the input state (colored) and in the output state (grey). The FNR was used for structural alignment. Proteins are shown in cartoon representation. The rotational axis of the FMND determined using Dyndom 3D is shown as a black line. Rotational movement of FMND is indicated by arrow. g, The structure of iNOS RED in the input state shown in cylinders. FMND is shown in solid and FNR is shown in transparency. FMN is colored green. Movement of FMND from the input state to the output state is denoted by an arrow. h, The structure of iNOS RED in the output state shown in cylinders. FMND is shown in solid and FNR is shown in transparency. FMN is colored green.

At the interface between the FNR and OXY, Y806 on the β11-β12 loop of the FNR inserts itself between R391 and R392 on α8 of the OXY (Fig. 4c), and K811 on the β12-α8 loop of the FNR forms an electrostatic interaction with E396 on the α8-α9 loop of the OXY (Fig. 4c). At the interface between the FMND and OXY, E540 on the β1-α1 loop of the FMND interacts with Q199 and N202 on the α3-β3 loop of the OXY (Fig. 4d), consistent with previous data showing that E762R and K423E mutations in nNOS (corresponding to E540 and N202 in iNOS) decrease NO production ^34,37,38^; E545 on α1 of the FMND forms an electrostatic interaction with K439 on α10 of the OXY (Fig. 4d), in line with previous results showing that the E545R mutation decreases NO formation of iNOS ^26^; K543 on α1 of the FMND interacts electrostatically with E435 on α10 of the OXY (Fig. 4d); N589 on the β3-α3 loop of the FMND interacts with N230 and N233 on the α4-β4 loop of the OXY (Fig. 4e); and Y625 on the β4-α4 loop of the FMND forms a hydrogen bond with T426 on the α9-α10 loop of the OXY (Fig. 4e). These two interfaces stabilize the docking of both the FNR and the FMND onto the OXY dimer and are dominated by hydrogen bonding and electrostatic interactions, consistent with previous data showing that the electron transfer efficiency from FMN to haem is regulated by ionic strength in a bell-shaped manner ^39^.

### Rotational movements of FMND during iNOS catalysis

We compared the structures of the RED between the input state and the output state (Fig. 4f-h). When the FNR is aligned, the FMND-CaM exhibits an outward rotation of up to 50°(Fig. 4d). This rotational movement of the FMND alternately positions the isoalloxazine group of FMN either toward FAD inside the RED (Fig. 4g) or toward the OXY bound to the exterior surface of the RED (Fig. 4h) for electron shuttling, highlighting the dynamic movements of the FMND during catalysis for electron shuttling.

### Mutation on the FMND-OXY interface traps iNOS in an intermediate state

During structural analysis, we identified S594 on the FMND as a residue located at the FMND-OXY interface in the output state (Fig. 4e), whereas it is not involved in the interaction between the FMND and FNR. The corresponding residue in nNOS is a glutamate (E816, Extended Data Fig. 8), and the activity of iNOS is higher than that of nNOS ^40^. Additionally, previous studies have shown that mutating E816 of nNOS to alanine, asparagine, or arginine decreases haem reduction rate and NO production activity ^34^, likely due to an altered interface between the FMND and OXY. Therefore, we hypothesized that the S594E mutation in iNOS might destabilize the FMND-OXY interface, thereby reducing its activity. Consistent with this hypothesis, the S594E mutation greatly decreased the enzymatic activity of iNOS (Fig. 5a). To investigate the structural basis of this activity loss, we determined the cryo-EM structure of the S594E mutant (Fig. 5b–c and Extended Data Fig. 9). During 2D classification, we observed particles representing the isolated OXY dimer, isolated RED, and particles showing both an OXY dimer and RED (Extended Data Fig. 9b). We focused on particles with both OXY and RED for further 3D classification and identified two major classes: Class 1 resembled the output state of wild-type iNOS (Extended Data Fig. 9c), whereas Class 2 exhibited a distinct RED conformation compared to the output state (Extended Data Fig. 9i–j). Although the final reconstruction of Class 2 reached only an overall resolution of 4.2 Å (Extended Data Fig. 9, and Supplementary Table 1)—insufficient for side-chain modeling—we could confidently dock each domain, together with their bound ligands into the density. Importantly, this conformation is neither the input state nor the output state; rather, the orientation of the FMND lies between the two states (Fig. 5d–h). In addition, the distances between FAD and FMN and between FMN and haem are too large to allow efficient electron transfer (Fig. 5i-j). This analysis suggests that a subset of particles in the S594E mutant is trapped in a non-productive intermediate conformation. Notably, E594 faces the solvent and does not form direct interactions with any other residues (Extended Data Fig. 9k), indicating that this conformation is not induced by new interactions introduced by the S594E mutation. Together, these structural observations provide a mechanistic explanation for the reduced activity of the S594E mutant: the mutation likely inhibits the conformational transition to the productive output state.

**Figure 5.**
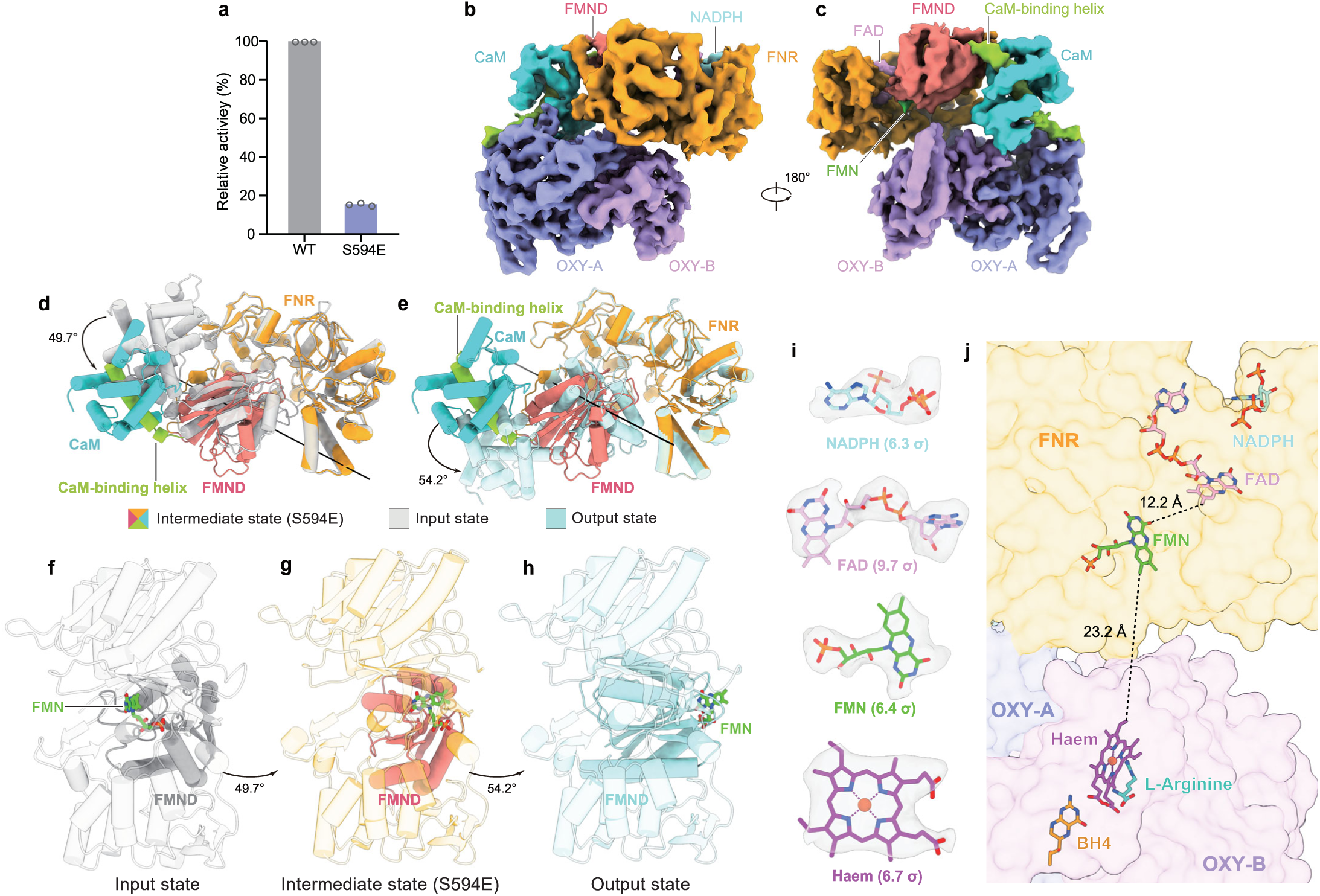
Structure of the iNOS S594E mutant. a, NO-producing activity of wild-type iNOS and the S594E mutant. Each data point is shown as an open circle. Data are mean ± s.d.; n = 3 technical replicates. The experiment was performed twice independently with similar results. WT, wild type. b, Cryo-EM map of the iNOS S594E mutant. Each subunit is colored as in Fig. 1b. c, A 180°-rotated side view of b. d, Structural comparison between the iNOS S594E mutant (colored) and iNOS in the input state (grey). The FNR was used for structural alignment. e, Structural comparison between the iNOS S594E mutant (colored) and iNOS in the output state (light blue). The FNR was used for structural alignment. f, The structure of iNOS RED in the input state shown in cylinders. FMND is shown in solid and FNR is shown in transparency. FMN is colored in green. Movement of FMND from the input state to the intermediate state is denoted by an arrow. g, The structure of iNOS RED in the intermediate state shown in cylinders. FMND is shown in solid and FNR is shown in transparency. FMN is colored in green. Movement of FMND from the intermediate state to the output state is denoted by an arrow. h, The structure of iNOS RED in the output state shown in cylinders. FMND is shown in solid and FNR is shown in transparency. FMN is colored in green. i, Densities of substrates and cofactors found in the iNOS-CaM complex in the intermediate state. Densities corresponding to NADPH, FAD, FMN, and Haem were contoured at 6.3 σ, 9.7 σ, 6.4 σ, and 6.7 σ, respectively. The nicotinamide group of NADPH is not modeled due to its poor density. j, Relative positions of cofactors and substrates in the intermediate state structure. Edge-to-edge distances between the isoalloxazine groups of FMN and FAD, as well as between FMN and haem, are denoted beside the dashed lines.

## Discussion

iNOS is a remarkably dynamic enzyme. In this work, we resolved the structures of two major functional states of wild-type iNOS captured during its catalytic cycle: the input state and the output state. It is reasonable to speculate that the iNOS holoenzyme undergoes continuous conformational transitions between these two states to perform catalysis. However, due to the low occupancy and intrinsic heterogeneity of intermediate conformations, we were unable to resolve them at high resolution. Additional approaches, such as chemical crosslinking or site-directed mutagenesis, may prove useful for trapping or enriching these transient states. Indeed, the introduction of the S594E mutation at the FMND-OXY interface not only decreased NO-producing activity but also enriched an intermediate conformation of iNOS (Fig. 5). For a significant portion of S594E particles, the FMND adopts a conformation that lies between the canonical input and output states, likely representing a transient snapshot captured prior to full engagement of the output state (Fig. 5d-h). By integrating these three structures, we constructed a hypothetical conformational trajectory of iNOS during catalysis (Extended Data Fig. 10 and Supplementary Video 1). In this model, iNOS transitions from the input state to an intermediate state—exemplified by the S594E mutant—and then reaches the output state to transfer electrons to haem, after which it returns to the input state to initiate another round of electron shuttling.

To accommodate the requisite dynamic motions, the interdomain interfaces of iNOS must undergo substantial reorganization. Specifically, the FMND–FNR and FMND–OXY interfaces must be sufficiently stable to ensure correct domain docking geometry for efficient electron transfer, yet sufficiently labile to permit dissociation and shuttling of the FMND. Interestingly, we found that highly conserved E540 and N589 of the FMND (Extended Data Fig. 7) participate in interactions with both the FNR in the input state (Fig. 2) and the OXY in the output state (Fig. 4), underscoring their functional importance during the motions of the FMND. Moreover, in addition to dynamic protein–protein interactions, the redox status of the FMN cofactor likely plays a pivotal role in modulating these dynamics. The FMND containing fully oxidized FMN may exhibit high affinity for the FNR but low affinity for the OXY. Upon reduction, the FMND harboring reduced FMN may display reduced affinity for the FNR and enhanced affinity for the OXY. This model is partly supported by both experimental data ^41^ and molecular dynamics simulations ^42^.

Our model also shows the detachment of the RED from the OXY dimer during electron shuttling in iNOS (Extended Data Fig. 10 and Supplementary Video 1). When we superpose the RED from the input state onto the output state structure by aligning their FNR, we observe multiple steric clashes between the FMND in its input-state conformation and the OXY dimer in the output state (Extended Data Fig. 5b). This analysis demonstrates that the input state is sterically incompatible with stable association with the OXY dimer, providing a mechanistic rationale for why the entire RED must dissociate from the OXY dimer to enable electron transfer from FAD to FMN—a critical prerequisite for recharging the FMN cofactor prior to the next output event.

Although CaM wraps tightly around the CaM-binding helix of iNOS (Fig. 4b), it exhibits limited interactions with either the FNR or OXY in the output state, which accounts for its relatively blurry electron density (Fig. 3b). A comparative analysis of our input and output state structures with the crystal structure of the isolated FMND–CaM complex (PDB ID: 3HR4 ^21^) reveals a wobbling movement of CaM relative to the RED (Extended Data Fig. 5c). This observation is consistent with the high mobility of CaM inferred from our cryo-EM maps. Notably, hydrogen–deuterium exchange experiments have suggested that CaM can directly interact with the OXY ^26^; therefore, it is plausible that CaM can transiently engage the OXY to facilitate the subsequent docking of the RED during iNOS catalysis. Furthermore, we identified steric clashes between CaM and the β11-β12 loop of the FNR in the context of the nNOS RED in the input state (PDB ID: 1TLL ^23^) (Extended Data Fig. 5a), suggesting that CaM binding may remodel the configuration of the RED to enhance the conformational dynamics of the FMND in nNOS. This model is consistent with previous biochemical and biophysical studies demonstrating that CaM binding increases FMND mobility and accelerates electron transfer from FMN to haem in NOS enzymes ^41,43–45^.

Sequence alignment across iNOS, nNOS, and eNOS reveals a high degree of conservation among residues that constitute the FMND-FNR and FMND–OXY interfaces (Extended Data Fig. 6-8), suggesting that nNOS and eNOS likely employ the same set of conserved residues as iNOS to mediate docking of the FMND onto the FNR or OXY dimer. In contrast, the FNR–OXY interface exhibits considerably lower sequence conservation (Extended Data Fig. 6 and 8). Additionally, the FNR–OXY interface undergoes reorganization in the S594E mutant structure (Extended Data Fig. 9i-j), further underscoring its dynamic nature. Therefore, whether the FNR of nNOS and eNOS engage the OXY dimer in the output state in a manner analogous to that observed in iNOS remains an open question that warrants further investigation.

Taken together, this work provides critical structural insights into the electron shuttling mechanism of iNOS and establishes a framework for understanding the dynamic conformational rearrangements that underpin NOS catalysis. Given the high degree of structural and mechanistic conservation among the three mammalian NOS isoforms, these findings have broad implications for our understanding of eNOS and nNOS function. Moreover, the domain-shuttling mechanism described here may also be relevant to other distantly related diflavin reductases and P450-like enzymes that employ modular domain motions to facilitate interdomain electron transfer ^46^.

## Methods

### Cell cultures

Sf9 insect cells (Thermo Fisher Scientific) were cultured in SIM SF (Sino Biological) at 27 °C. Adapted Expi293F cells (donated by Shanghai OPM Biosciences) were cultured in 293F Hi-exp Medium (Shanghai OPM Biosciences) at 37 °C with 6% CO2 and 70% humidity. The cell lines were routinely checked to be negative for mycoplasma contamination but have not been authenticated.

### Constructs

To express the wild-type mouse iNOS-CaM complex, the DNA sequence encoding wild-type mouse iNOS was cloned into the pBMCL1 vector ^33^ with the sequence of green fluorescent protein (GFP) and tandem affinity tags (His-Strep-Flag-Strep) fused to its C-terminus, thus generating pBMCL1-iNOS-CGFP-Ctag. The cDNA of iNOS and GFP were flanked by the sequence of the HRV3C site for the removal of GFP and tags before size-exclusion chromatography (SEC). The coding sequence of mouse CaM was also cloned into the pBMCL1 vector with the ALFA tag, mScarlet ^47^, and HRV3C protease site introduced at its N-terminus, generating pBMCL1-ALFA-mScarlet-CaM. The expression vector of the iNOS S594E mutant was generated based on pBMCL1-iNOS-CGFP-Ctag using the QuikChange method.

### Protein expression and purification

Baculoviruses of iNOS-CGFP-Ctag, iNOS (S594E)-CGFP-Ctag, and ALFA-mScarlet-CaM were generated using Sf9 cells for protein expression. Adapted Expi293F cells were co-infected with P2 viruses of iNOS and CaM (10% volume, iNOS: CaM = 8: 2) at a density of 3.0 × 10^6^ cells per ml. The cell culture was supplemented with sodium butyrate (10 mM) to boost protein expression and 5-aminolevulinic acid hydrochloride (200 μM) to promote haem incorporation 12 h after the addition of P2 viruses, and the cells were transferred to a 30 °C incubator for another 36 h before collection. The cells were collected by centrifugation at 4,000g (JLA-8.1000, Beckman Coulter) for 10 min at 4 °C, washed with TBS buffer (20 mM Tris pH 8.0 at 4 °C and 150 mM NaCl) containing 2 μg ml^−1^ aprotinin, 2 μg ml^−1^ pepstatin, and 2 μg ml^−1^ leupeptin, then flash-frozen in liquid nitrogen and stored at −80 °C.

For protein purification, the cell pellet was resuspended in lysis buffer (50 mM Tris pH 8.0 at 4 °C, 150 mM NaCl, 2 μg ml^−1^ aprotinin, 2 μg ml^−1^ pepstatin, 2 μg ml^−1^ leupeptin, 20% glycerol (v/v), 1 mM CaCl_2_, 1 mM phenylmethanesulfonyl fluoride (PMSF), 10 μM BH4, 0.5 mM Tris(2-carboxyethyl)phosphine (TCEP), 10 mM MgCl_2_, and 10 μg ml^−1^ DNAse). The mixture was sonicated for 20 min at 4 °C, and insoluble debris was removed by centrifugation at 193,400g (Ti50.2, Beckman Coulter) for 40 min. The supernatant was then loaded onto a 3 ml Streptactin Beads 4FF (Smart Lifesciences) column to purify the iNOS-CaM complex. The column was washed with 20 ml buffer A (20 mM Tris pH 8.0 at 4 °C, 150 mM NaCl, 10 μM BH4, 0.5 mM TCEP), followed by 50 ml buffer 1 (buffer A supplemented with 350 mM NaCl) to wash off nucleic acids, and 50 ml buffer 2 (buffer A supplemented with 10 mM MgCl2 and 2 mM ATP) to remove heat-shock protein impurities. The column was then extensively washed with 30 ml buffer A, and the target protein was eluted with buffer A supplemented with 5 mM d-desthiobiotin (IBA). A Soret peak at 400 nm was observed in the Streptactin eluate, suggesting the presence of BH4-bound functional iNOS ^48^. The iNOS concentration was calculated according to its absorbance at 400 nm, and GST-tagged HRV3C Protease was added to the eluate and incubated overnight to cleave the fluorescent tags on iNOS and CaM. The GST-HRV3C Protease was removed after cleavage using 200 μl GST resin (GE Healthcare). The GST flow-through was concentrated and loaded onto a Superose 6 increase 10/300 GL column (GE Healthcare) in S6 buffer (20 mM HEPES pH 7.5, 75 mM NaCl, and 10 μM BH4). Peak fractions corresponding to the dimeric iNOS-CaM complex were combined and concentrated for cryo-EM sample preparation. Both the wild-type iNOS-CaM and iNOS (S594E)-CaM complexes were obtained using the same workflow.

### Enzymatic assays

The NO-generating activities of iNOS were measured using the oxyhemoglobin assay ^49^ with minor modifications. Briefly, the change in NO concentration in the reaction mix was reflected by the oxidation of oxyhemoglobin to methemoglobin using an approximated extinction coefficient (Δ_ε401_) of 60,000 M^−1^cm^−1^. A 100 μl reaction mix contained 20mM Tris (pH 7.5 at room temperature), 75 mM NaCl, 10 μM BH4, 100 U superoxide dismutase (SOD), 50 U catalase, 100 μM NADPH, 500 μM Arg, 6 μM HbO2, 1 μM FAD, 1 μM FMN, and 10 nM iNOS protein. The reaction was initiated by adding purified iNOS to the reaction mix containing the other reagents and immediately subjected to UV-Vis absorbance measurement at 401 nm using a Microplate Reader (Tecan Infinite M Plex) at 30 °C. The A401-to-time curve (see Extended Data Fig.1a) reflected the NO-generating activity of the iNOS sample. The relative activity of iNOS mutants was normalized to that of wild-type iNOS (see Fig.5a).

### Cryo-EM sample preparation and data collection

Both the wild-type iNOS-CaM complex and the iNOS (S594E)-CaM complex were concentrated to approximately 4.7 μM with an A280 of 3.5. FAD, FMN, and L-Arginine were added to the protein sample to final concentrations of 0.5 mM, 0.5 mM, and 2 mM, respectively. The protein mix was ultracentrifuged for 30 min, and the supernatant was further supplemented with 2 mM NADPH to initiate the catalytic reaction, which was incubated at 37 ℃ for 30 s to reach turnover conditions. Then, 0.5 mM fluorinated octyl maltoside (FOM) was added to the sample to improve orientation distribution. Aliquots of 2.5 μl sample were quickly applied to pre-discharged Quantifoil Au 300 mesh R 0.6/1.0 grids. After a 5-s incubation on the grids’ surface at room temperature under 100% humidity, the grids were blotted for 3.5 s using a blot force of 2, then plunge-frozen into liquid ethane using a Vitrobot Mark IV (Thermo Fisher Scientific). Cryo-EM grids were screened on a Talos Arctica electron microscope (Thermo Fisher Scientific) operating at 200 kV using a K2 camera (Thermo Fisher Scientific), and selected grids were transferred to a Titan Krios electron microscope (Thermo Fisher Scientific) operating at 300 kV for data acquisition. EPU-2.12.1.2782REL was used for automated data collection. Images were collected with a K3 Summit direct electron detector (Gatan) mounted after a quantum energy filter (slit width 20 eV) in super-resolution mode with a defocus range of −1.5 μm to −1.8 μm.

Data for the wild-type iNOS-CaM complex were collected at a magnification of 60,000×, resulting in a calibrated super-resolution pixel size of 0.417 Å. The defocus ranged from −1.5 to −1.8 μm, with a total dose of 50 e−/Å^2^ and a dose rate of 22.15 e^−^/pixel/s on the detector.

Data for the iNOS (S594E)-CaM complex were collected at a magnification of 47,000× with a super-resolution pixel size of 0.534 Å, and the defocus ranged from −1.5 to −1.8 μm, with a total dose of 50 e−/Å^2^ and a dose rate of 22.36 e^−^/pixel/s on the detector.

### Cryo-EM data processing

A total of 2,731 multi-frame movies of the wild-type iNOS-CaM complex under turnover conditions were collected. Beam-induced motion was corrected using MotionCor2 ^50^ and binned to a pixel size of 0.834 Å. The data processing workflow was performed in cryoSPARC ^51^. Contrast transfer function (CTF) parameters of dose-weighted micrographs were estimated using the patch CTF function. A total of 2,160,734 particles were template-picked using the previously resolved density of the iNOS OXY dimer (data not shown) as the reference, binned, and extracted with a box size of 150 pixels. Two-dimensional (2D) classification and 3D classification were performed to remove ice, contaminants, aggregates, and particles with unrecognizable features, yielding 47,984, 542,442, and 37,894 particles for the input state (RED alone), OXY alone, and the output state, respectively. Particles corresponding to the input state and output state were further classified using methods including 3D variability analysis (3DVA)^52^, multi-reference heterogeneous refinement, 3D classification without alignment, and HR-HAIR high-resolution reconstruction ^53^. The highly purified particles representing the input and output states were used as references for Topaz picking ^54^ and seeds for seed-facilitated classification ^55^, resulting in two enlarged particle sets. Particles representing the output state were re-extracted with a box size of 300 pixels, followed by multi-reference 3D classification, resolution-gradient 3D classification, and non-uniform refinement (NU-refinement) to further improve map quality, obtaining consensus maps at 3.82 Å. To further improve local map quality, the particles were subjected to signal subtraction using the masks shown in Extended Data Fig. 2 and local refinement to generate focus-refined maps. For the output state, the resolutions of refined maps focused on OXY-CaM-FMND and CaM-FMND-FNR were 3.32 Å and 3.91 Å, respectively. These two locally-refined maps were aligned to the consensus map and merged using the UCSF Chimera ^56^ “vop maximum” command to generate a composite map for model building and interpretation. For the input state particle set, the CaM-FMND-FNR was further refined using a similar strategy to reach a resolution of 3.77 Å (see Extended Data Fig.2). Resolutions were estimated using the gold-standard Fourier shell correlation 0.143 criterion. Local-resolution maps were calculated using cryoSPARC.

A total of 1,126 multi-frame movies of the iNOS (S594E)-CaM complex under turnover conditions were collected. Data for the iNOS (S594E)-CaM complex were processed in a similar manner as shown in Extended Data Fig. 9.

### Model building, refinement, and validation

AlphaFold-3 (AF3) ^57^ predicted structures of the separated domains of the mouse iNOS-CaM complex (OXY dimer, CaM-CaM-binding helix, FMND, and FNR) were docked into the final maps in Chimera ^56^ and manually rebuilt using Coot ^58^. The resulting models were refined against the maps using PHENIX ^59,60^.

### Quantification and statistical analysis

Data processing and statistical analysis were conducted using GraphPad Prism software. Statistical details can be found in the methods and figure legends. Global resolution estimations of cryo-EM density maps are based on the Fourier shell correlation 0.143 criterion ^61^. Local resolution was estimated using cryoSPARC. The number of independent experiments (n) and relevant statistical parameters for each experiment (such as mean and s.d.) are described in the figure legends. No statistical methods were used to predetermine sample sizes.

## Data Availability

The data that support the findings of this study are available from the corresponding author upon request. Cryo-EM maps and atomic coordinates of iNOS in the input state have been deposited in the EMDB and PDB databases under the ID codes EMDB: EMD-80277 and PDB: 25OT, respectively. Cryo-EM maps and atomic coordinates of iNOS in the output state have been deposited in the EMDB and PDB databases under the ID codes EMDB: EMD-80251 and PDB: 25OI, respectively. Cryo-EM maps and atomic coordinates of the iNOS S594E mutant have been deposited in the EMDB and PDB databases under the ID codes EMDB: EMD-80278 and PDB: 25OU, respectively.

## Acknowledgments

We thank all members of the Chen Lab for their kind help. Cryo-EM data collection was supported by the Electron Microscopy Laboratory and the Cryo-EM platform of Peking University. Part of the structural computation was also performed on the Computing Platform of the Center for Life Science and the High-performance Computing Platform of Peking University. We thank the National Center for Protein Sciences at Peking University in Beijing, China, for assistance with negative stain EM. This work is supported by grants from the Ministry of Science and Technology of China (National Key R&D Program of China, 2022YFA1303000 to L.C.), the National Natural Science Foundation of China (32225027 to L.C.), and the Center for Life Sciences (CLS to L.C.).

## Author contributions

L.C. initiated the project. X.L. generated prokaryotic expression constructs and performed early work on NOS. Y.S. purified the protein, prepared the cryo-EM sample, collected the cryo-EM data, and built the model. All authors contributed to manuscript preparation.

---

**Extended Data Figure 1.**
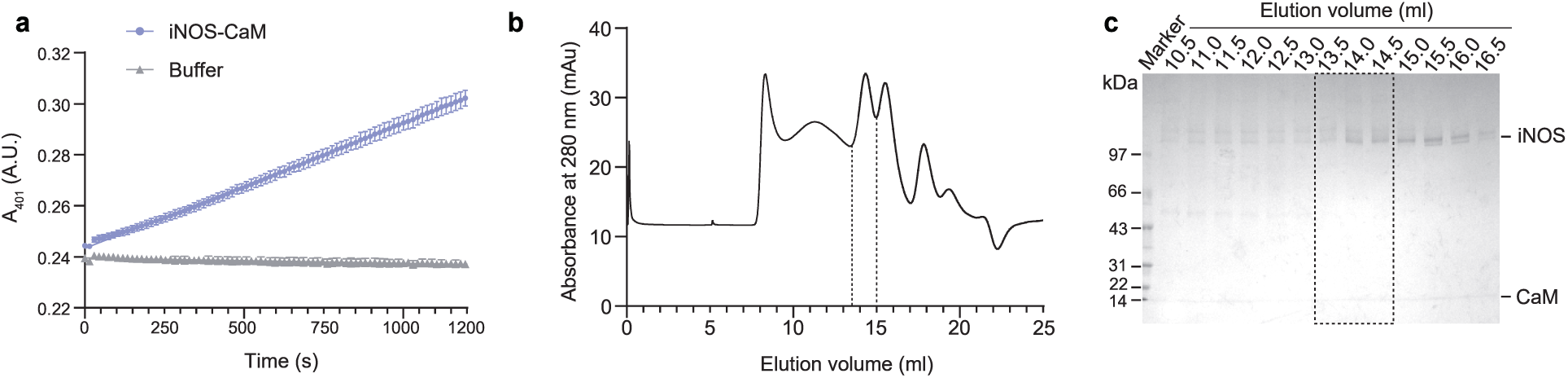
Protein expression and purification of iNOS. a, The NO-producing activity of the mouse iNOS-CaM complex measured by the oxyhemoglobin assay. The activity of the iNOS-CaM complex in the streptactin-affinity-chromatography eluate (see Methods, Protein expression and purification) and the corresponding elution buffer are represented by purple and gray lines, respectively. Data are mean ± s.d.; n = 3 technical replicates, and the experiment was repeated independently twice with similar results. b, Size-exclusion chromatography (SEC) profile of the wild-type iNOS-CaM complex. Fractions between the dashed lines represent the assembled iNOS dimer and were used for cryo-EM sample preparation. c, Coomassie brilliant blue-stained SDS–PAGE gel of the fractions eluted after size-exclusion chromatography. Lanes boxed by dashed lines represent the fractions used for cryo-EM sample preparation, corresponding to the pooled fractions in (b).

**Extended Data Figure 2.**
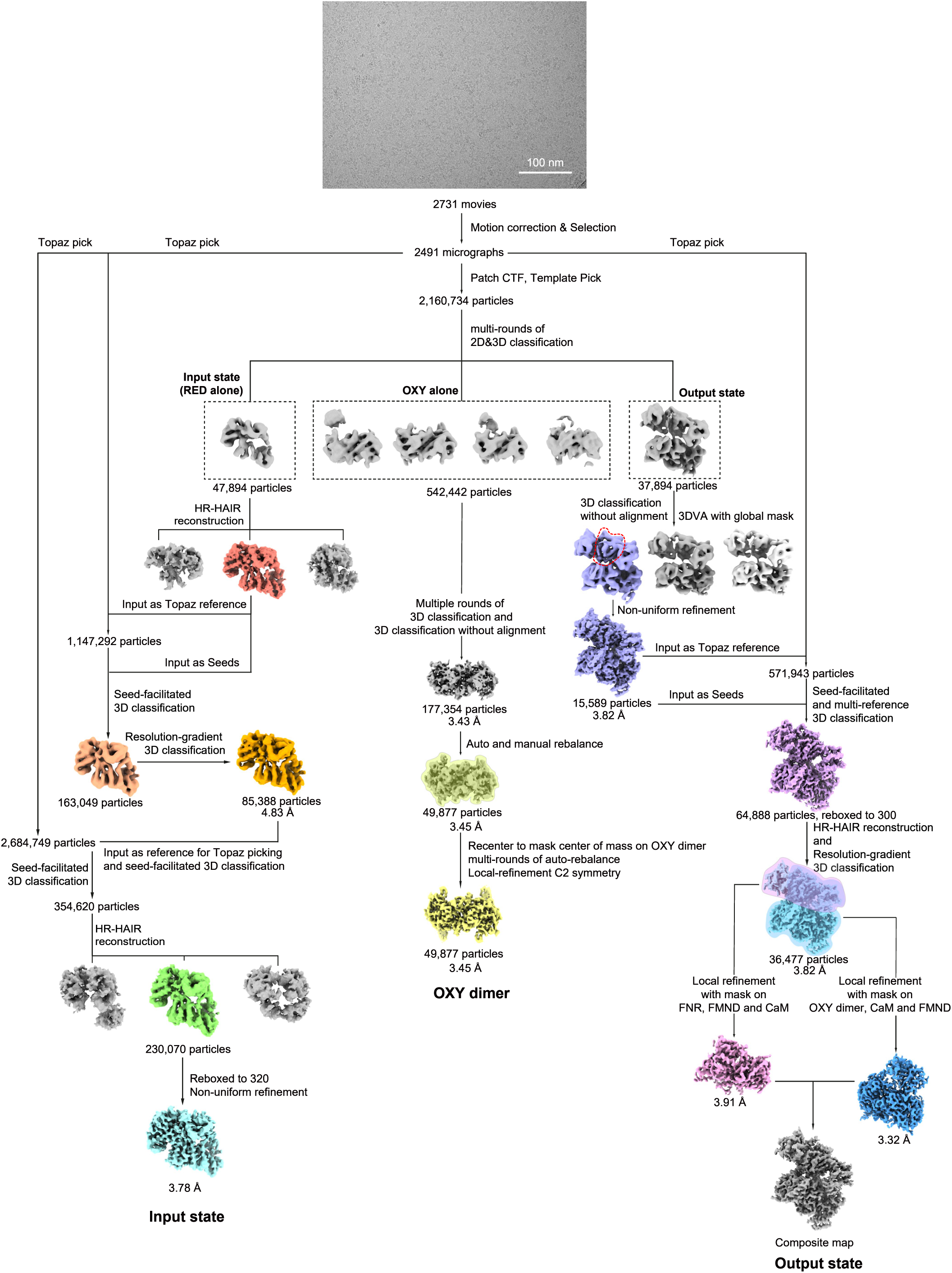
Cryo-EM image processing of wild-type iNOS under turnover conditions. A representative raw micrograph (2,731 in total) of the wild-type iNOS-CaM complex under turnover conditions is shown at the top. The image processing workflow is shown below. The processing procedures for the input state (RED alone), OXY alone, and output state classes are shown on the left, in the middle, and on the right, respectively.

**Extended Data Figure 3.**
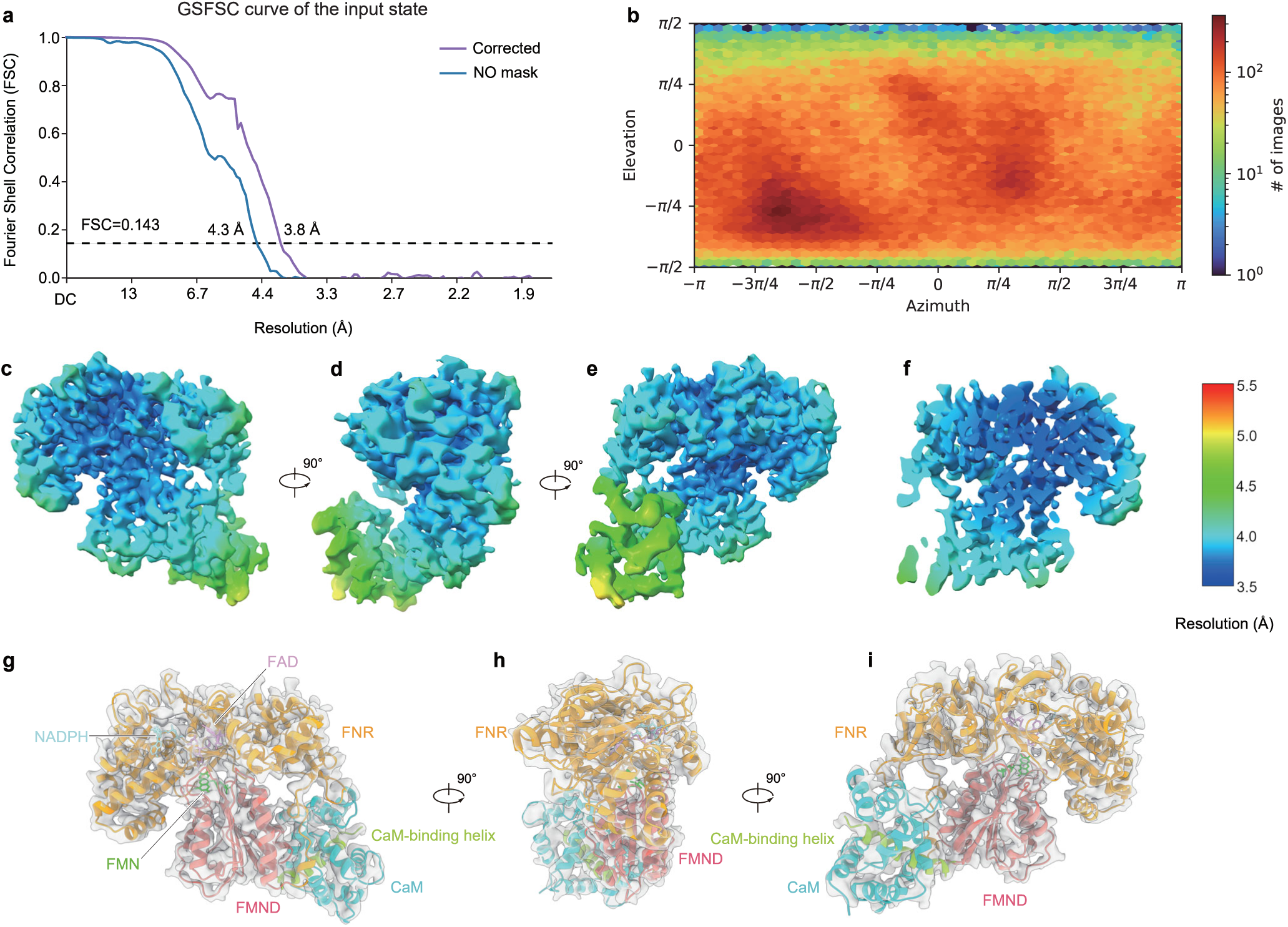
Cryo-EM map of wild-type iNOS in the input state. a, Gold-standard Fourier shell correlation (GSFSC) curves of the consensus refinement of wild-type iNOS in the input state. b, Angular distributions of the consensus refinement of wild-type iNOS in the input state. c, Local resolution distribution of wild-type iNOS in the input state after consensus refinement. d, A 90°-rotated view of (c). e, A 90°-rotated view of (d). f, A cut-open view of (e). g, Fitting of RED-CaM complex into the focus-refined cryo-EM map. h, A 90°-rotated view of (g). i, A 90°-rotated view of (h).

**Extended Data Figure 4.**
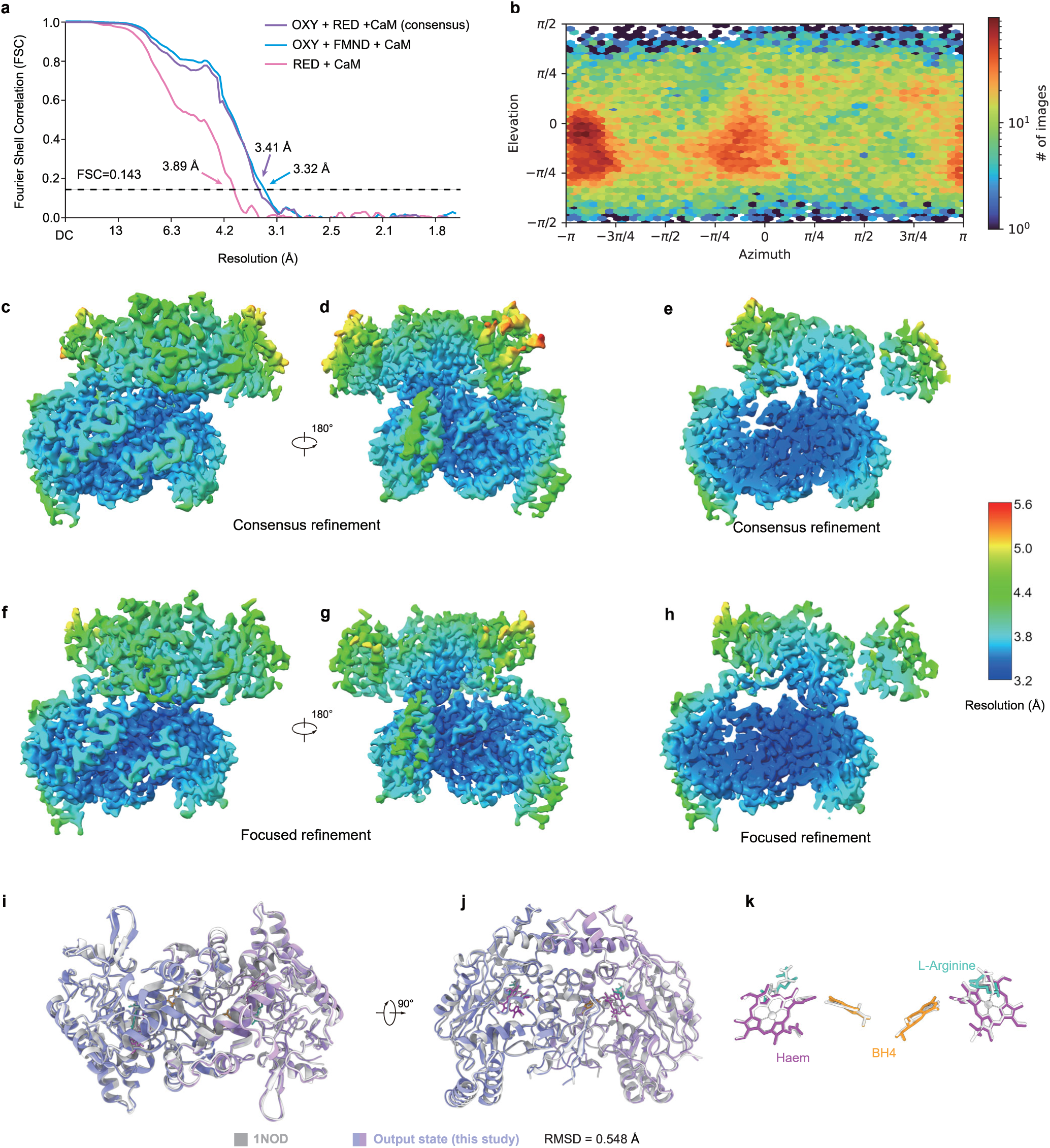
Cryo-EM map of wild-type iNOS in the output state. a, Gold-standard Fourier shell correlation (GSFSC) curves of the consensus refinement of wild-type iNOS in the output state. b, Angular distributions of the consensus refinement of wild-type iNOS in the output state. c, Local resolution distribution of wild-type iNOS in the output state after consensus refinement. d, A 180°-rotated view of (c). e, A cut-open view of (c). f, Local resolution distribution of wild-type iNOS in the output state after focused refinement. g, A 180°-rotated view of (f). h, A cut-open view of (f). i, Structural comparison between the OXY region of the iNOS-CaM complex in the output state (this study, colored as in Fig. 1) and the previously reported crystal structure of the mouse iNOS OXY dimer (PDB: 1NOD, colored white). j, A 90°-rotated view of (i). k, A zoom-in view of the ligands in (j).

**Extended Data Figure 5.**
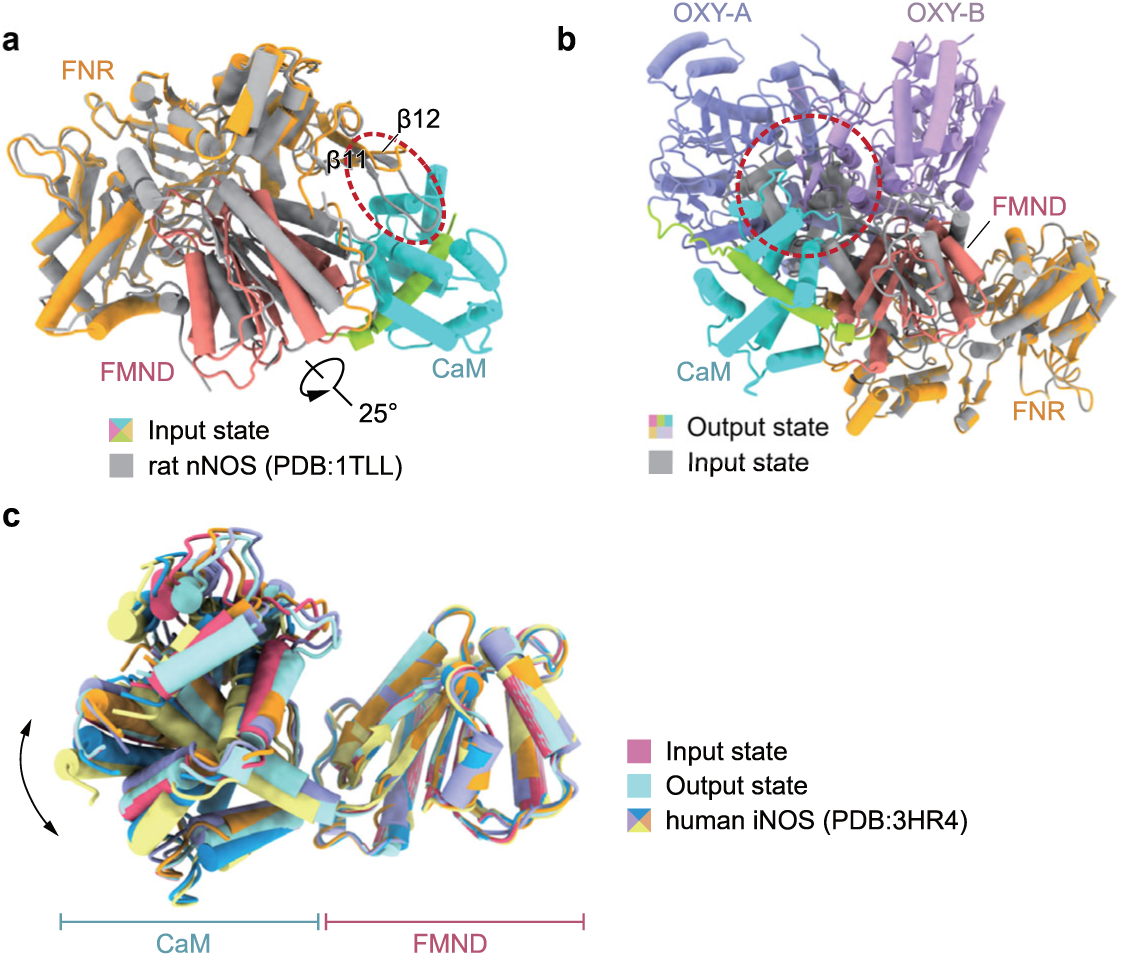
Structure comparisons. a, Structural comparison between the mouse iNOS structure in the input state (colored) and the crystal structure of rat nNOS RED (gray, PDB ID: 1TLL). The FNR was used for structural alignment. The rotation of the FMND is indicated by an arrow. Steric clashes between CaM in the iNOS structure and the FNR of nNOS are indicated with a red circle. b, Structural comparison between iNOS in the output state (colored) and iNOS in the input state (gray). The FNR was used for structural alignment. Steric clashes between the FNR of the input state and the OXY in the output state are indicated with a red circle. c, Structural comparison between iNOS in the input state (pink), in the output state (light blue), and crystal structures of the human iNOS FMND-CaM complex in four conformations (colored, PDB ID: 3HR4). The FMND was used for structural alignment. Rotations of CaM are indicated by arrows.

**Extended Data Figure 6.**
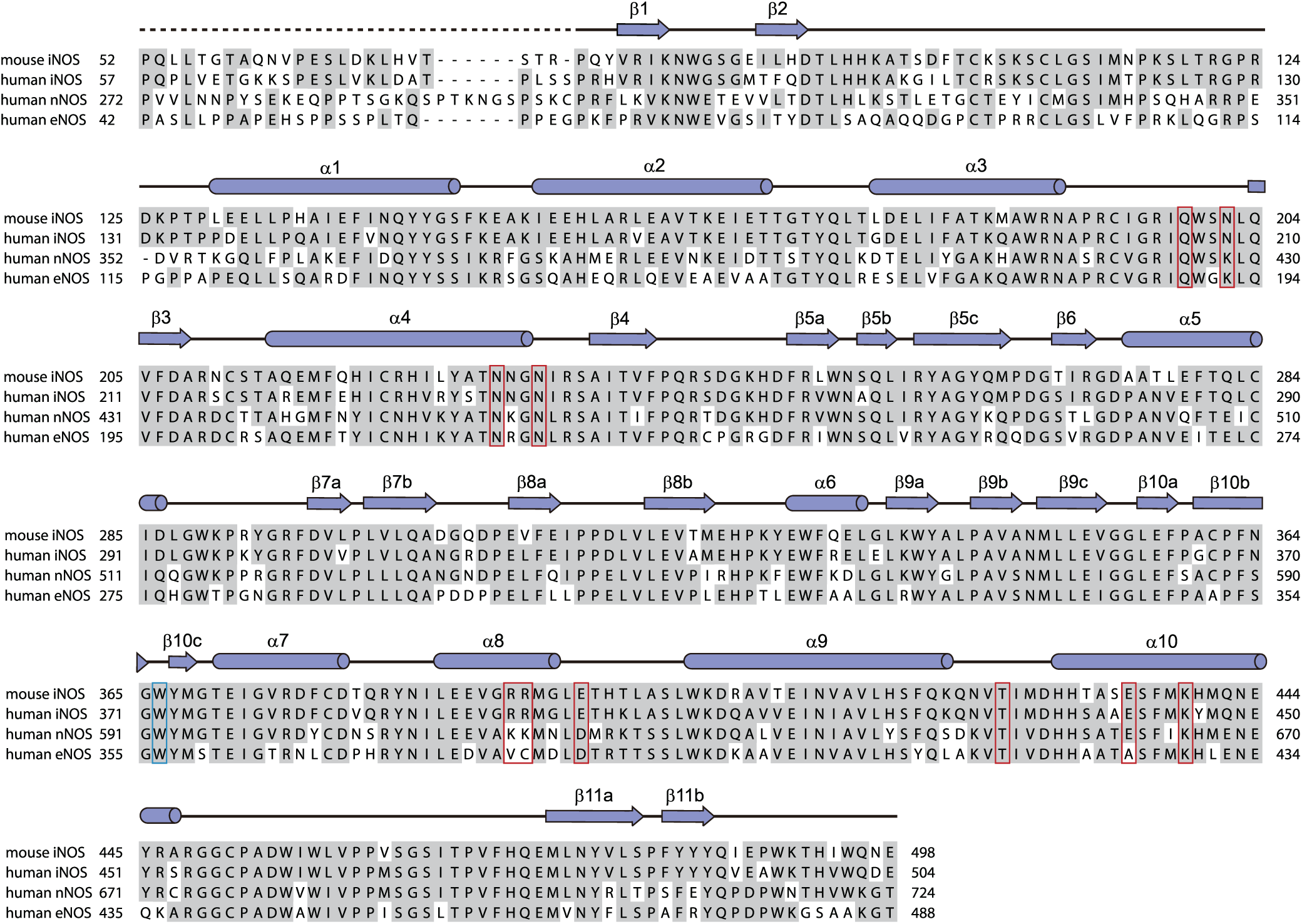
Sequence alignment of the oxygenase domain (OXY) Multiple sequence alignment (MSA) of the OXY domain from Mus musculus iNOS (UniProt ID: P29477), Homo sapiens iNOS (UniProt ID: P35228), Homo sapiens nNOS (UniProt ID: P29475), and Homo sapiens eNOS (UniProt ID: P29474). Sequences were downloaded from UniProt and aligned using ClustalX. The MSA result was further processed in BioEdit. Conserved residues are highlighted in gray. Secondary structures are indicated as cylinders (α helices), arrows (β sheets), and lines (loops). Unmodeled residues are indicated by dashed lines. W366 near the haem center is boxed in blue. Residues that interact with the FNR and FMND are boxed in red.

**Extended Data Figure 7.**
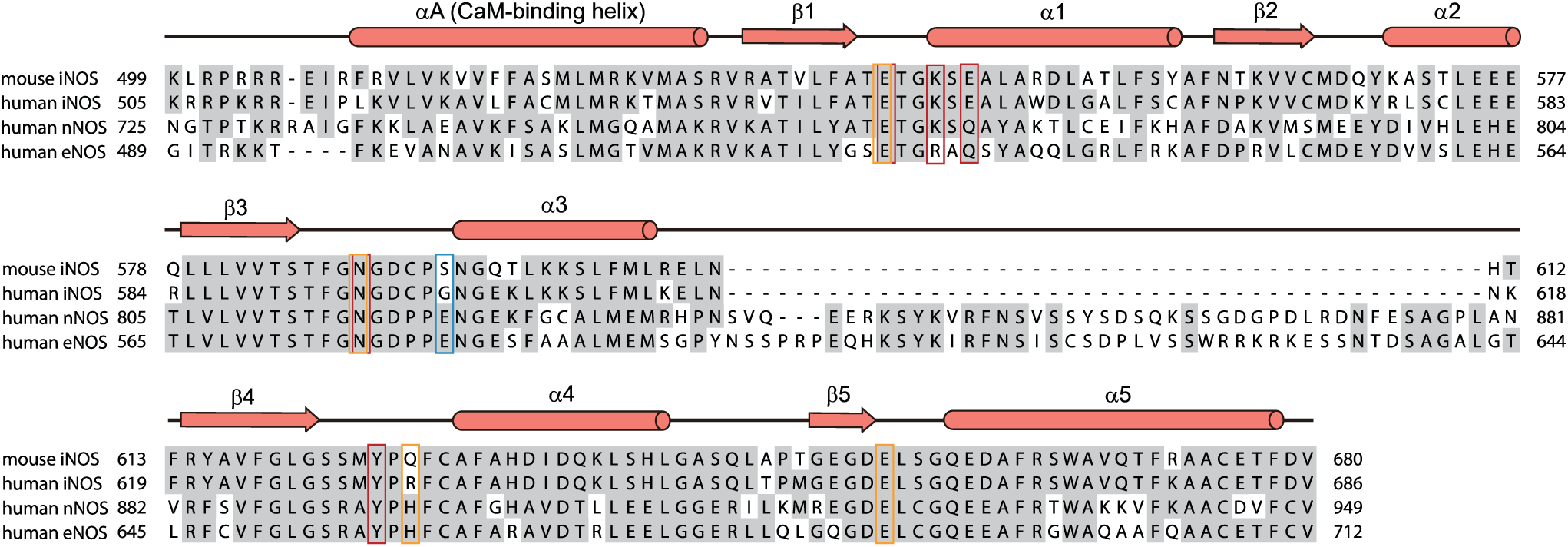
Sequence alignment of the FMN-binding subdomain (FMND) Multiple sequence alignment (MSA) of the FMND from Mus musculus iNOS (UniProt ID: P29477), Homo sapiens iNOS (UniProt ID: P35228), Homo sapiens nNOS (UniProt ID: P29475), and Homo sapiens eNOS (UniProt ID: P29474). Sequences were downloaded from UniProt and aligned using ClustalX. The MSA result was further processed in BioEdit. Conserved residues are highlighted in gray. Secondary structures are indicated as cylinders (α helices), arrows (β sheets), and lines (loops). S594, residues interacting with OXY and residues interacting with FNR are boxed in blue, red and orange, respectively.

**Extended Data Figure 8.**
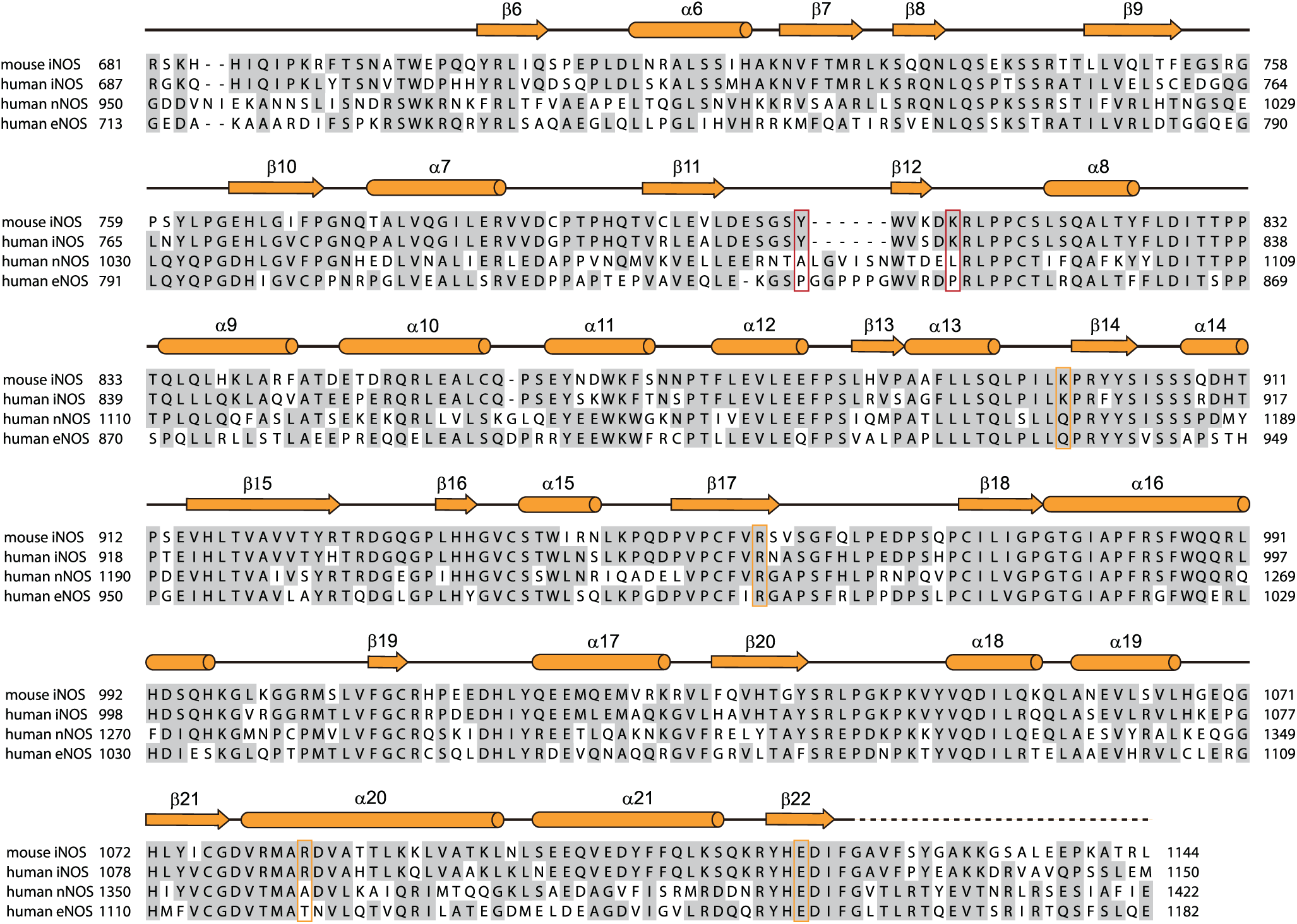
Sequence alignment of the FNR-like domain (FNR) Multiple sequence alignment (MSA) of the FNR from Mus musculus iNOS (UniProt ID: P29477), Homo sapiens iNOS (UniProt ID: P35228), Homo sapiens nNOS (UniProt ID: P29475), and Homo sapiens eNOS (UniProt ID: P29474). Sequences were downloaded from UniProt and aligned using ClustalX. The MSA result was further processed in BioEdit. Conserved residues are highlighted in gray. Secondary structures are indicated as cylinders (α helices), arrows (β sheets), and lines (loops). Residues interacting with OXY and FNR are boxed in red and orange, respectively.

**Extended Data Figure 9.**
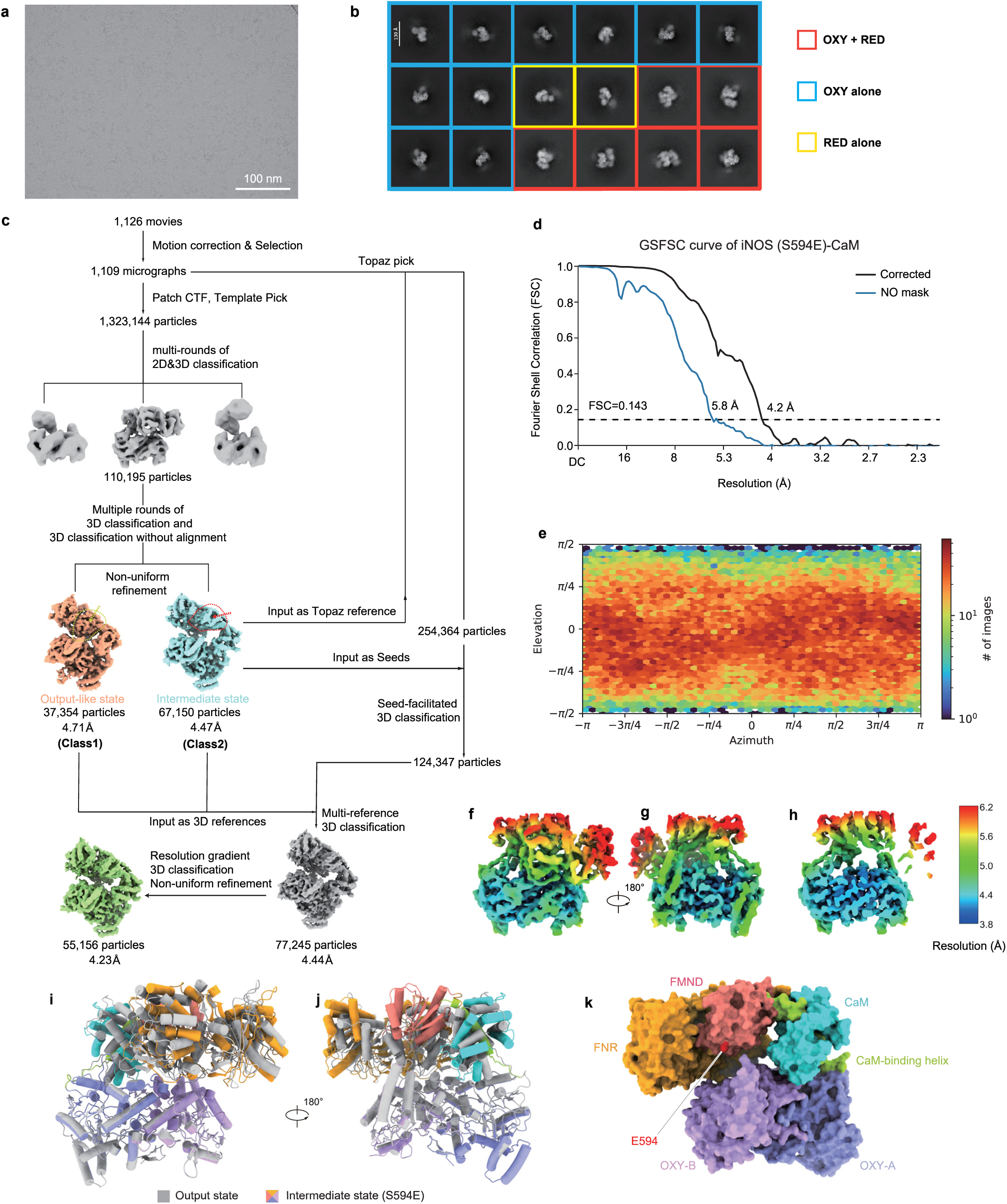
Cryo-EM image processing of the iNOS S594E mutant. a, A representative raw micrograph (1,126 in total) of the iNOS S594E mutant under turnover conditions. b, Representative 2D class averages of the iNOS S594E mutant. c, The cryo-EM data processing workflow of the iNOS S594E mutant. Differences in FMND orientation between the output-like state and the intermediate state are indicated by dashed lines and arrows. d, Gold-standard Fourier shell correlation (GSFSC) curves of the consensus refinement of the iNOS S594E mutant in the intermediate state. e, Angular distributions of the consensus refinement of the iNOS S594E mutant in the intermediate state. f, Local resolution distribution of the iNOS S594E mutant in the intermediate state after consensus refinement. g, A 180°-rotated view of (f). h, A cut-open view of (f). i, Structural comparison between wild-type iNOS in the output state (colored gray) and the iNOS S594E mutant in the intermediate state (colored as in Fig. 1b). j, A 180°-rotated view of (i). k, Structure of the iNOS S594E mutant in the intermediate state shown as a surface, colored as in Fig. 1. E594 is shown as spheres, colored red.

**Extended Data Figure 10.**
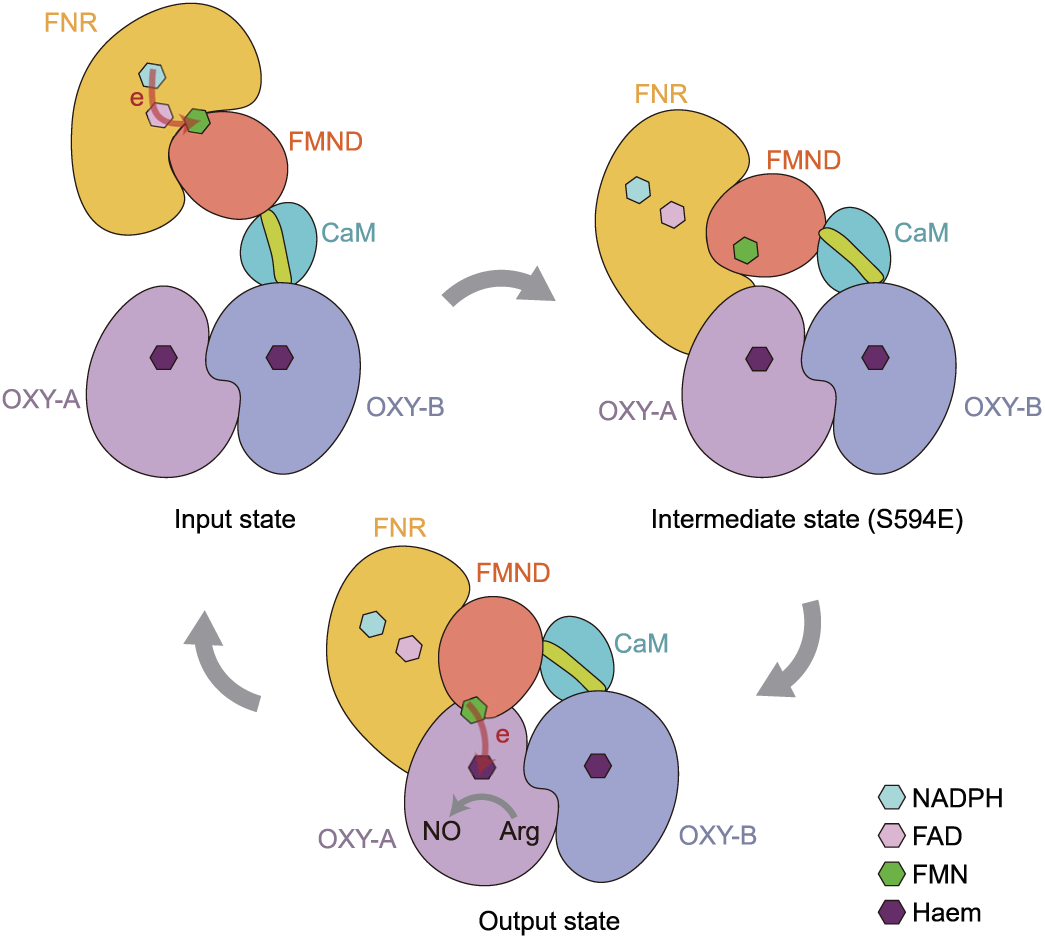
Model for electron shuttling in iNOS. Cartoon model of the conformational changes during iNOS catalysis. Each subunit is colored the same as in Fig. 1b. FMN, FAD, NADPH, and haem are denoted as hexagons. The electron transfer pathway is indicated with red arrows. In the input state, the FMND docks onto the FNR, allowing electrons to transfer from NADPH to FMN via FAD. In the intermediate state, the FMND rotates closer to the OXY, preparing for electron transfer to the OXY. In the output state, the FMND docks onto the OXY, allowing electrons to transfer from FMN to the haem center and subsequent production of nitric oxide. Transitions among these conformations enable efficient electron shuttling between subdomains.

**Supplementary Video 1. Conformational transitions of iNOS during catalysis** In this video, iNOS transitions from the input state to the intermediate state, then to the output state, and finally returns to the input state, completing one electron shuttling cycle. Three structures of iNOS—the input state, the intermediate state (S594E mutant), and the output state—were used as references for morph generation using Chimera X^62^.

**Supplementary Table S1.** Cryo-EM data collection, refinement and validation statistics

| PDB ID<br>EMDB ID | WT output<br>25OI<br>EMD-80251 | WT input<br>25OT<br>EMD-80277 | S594E<br>25OU<br>EMD-80278 |
| --- | --- | --- | --- |
| **Data collection and processing** |  |  |  |
| Magnification | 60,000 × |  | 47,000 × |
| Voltage (kV) | 300 |  | 300 |
| Electron exposure (e<sup>-</sup>/Å<sup>2</sup>) | 50 |  | 50 |
| Defocus range (μm) | -1.5 to -1.8 |  | -1.5 to -1.8 |
| Pixel size (Å) | 0.833 |  | 1.067 |
| Symmetry imposed | *C1* |  | *C1* |
| Initial particle images (no.) | 2,160,734 |  | 1,323,144 |
| Final particle images (no.) | 36,477 | 230,070 | 55,156 |
| Map resolution (Å) | 3.41<sup>a</sup> / 3.32<sup>b</sup> / 3.89<sup>c</sup> | 3.78 | 4.23 |
| FSC threshold | 0.143 | 0.143 | 0.143 |
| Map resolution range (Å) | 250-3.41 | 267-3.77 | 320-4.23 |
| **Refinement** |  |  |  |
| Initial model used (PDB code) | AF3 predicted | AF3 predicted | AF3 predicted |
| Model resolution (Å) | 3.5 | 3.9 | 4.5 |
| FSC threshold | 0.5 | 0.5 | 0.5 |
| Model resolution range (Å) | 250-3.5 | 267-3.9 | 320-4.5 |
| Map sharpening *B* factor (Å<sup>2</sup>) | -110.1<sup>a</sup> / -103.8<sup>b</sup> / -150.9<sup>c</sup> | -217.9 | -192.9 |
| **Model composition** |  |  |  |
| Non-hydrogen atoms | 13,179 | 6,137 | 13,051 |
| Protein residues | 1,614 | 765 | 1,610 |
| Ligands | 12 | 7 | 12 |
| ***B* factors (Å<sup>2</sup>)** |  |  |  |
| Protein | 204.71 | 165.17 | 204.71 |
| Ligand | 198.10 | 165.78 | 198.10 |
| **R.m.s. deviations** |  |  |  |
| Bond lengths (Å) | 0.003 | 0.004 | 0.004 |
| Bond angles (°) | 0.469 | 0.978 | 0.907 |
| **Validation** |  |  |  |
| MolProbity score | 1.56 | 1.63 | 1.52 |
| Clashscore | 9.00 | 10.94 | 8.34 |
| Poor rotamers (%) | 1.29 | 0.00 | 1.24 |
| **Ramachandran plot** |  |  |  |
| Favored (%) | 98.01 | 97.63 | 98.00 |
| Allowed (%) | 1.99 | 2.37 | 2.00 |
| Disallowed (%) | 0.00 | 0.00 | 0.00 |

